# Moderate adolescent chronic intermittent ethanol exposure sex-dependently disrupts synaptic transmission and kappa opioid receptor function in the basolateral amygdala of adult rats

**DOI:** 10.1101/2020.06.22.165811

**Authors:** Kathryn R. Przybysz, Meredith E. Gamble, Marvin R. Diaz

## Abstract

Adolescent alcohol exposure is associated with many negative outcomes that persist into adulthood, including altered affective and reward-related behaviors. However, the long-term neurological disruptions underlying these behavioral states are not fully understood. The basolateral amygdala (BLA) plays a critical role in many of these behaviors, and shifts in the excitatory/inhibitory balance in this area are capable of directly modulating their expression. While changes to BLA physiology have been demonstrated during the acute withdrawal phase following adolescent ethanol exposure, no studies to date have examined whether these persist long-term. The kappa opioid receptor (KOR) system is a neuromodulatory system that acts as a prominent mediator of negative affective behaviors, and alterations of this system have been implicated in the behavioral profile caused by chronic alcohol exposure in adulthood. Notably, in the BLA, the KOR system undergoes functional changes between adolescence and adulthood, but whether BLA KORs are functionally disrupted by adolescent ethanol exposure has not been examined. In this study, we exposed male and female Sprague-Dawley rats to a vapor inhalation model of moderate adolescent chronic intermittent ethanol (aCIE) and examined the long-term effects on GABAergic and glutamatergic neurotransmission within the adult BLA using whole-cell patch-clamp electrophysiology. We also assessed how KOR activation modulated these neurotransmitter systems in aCIE versus control rats using the selective KOR agonist, U69593. This investigation revealed that aCIE exposure disrupted basal glutamate transmission in females by increasing spontaneous excitatory postsynaptic current (sEPSC) frequency, while having no effects on glutamate transmission in males or GABA transmission in either sex. Interestingly, we also found that aCIE exposure unmasked a KOR-mediated suppression of spontaneous inhibitory postsynaptic current (sIPSC) frequency and sEPSC amplitude only in males, with no effects of aCIE exposure on KOR function in females. Together, these data suggest that moderate-level adolescent ethanol exposure produces long-term changes to BLA physiology and BLA KOR function, and that these changes are sex-dependent. This is the first study to examine persistent adaptations to both BLA physiology and KOR function following adolescent alcohol exposure, and opens a broad avenue for future investigation into other neurobiological and behavioral consequences of adolescent ethanol exposure-induced disruptions of these systems.

## Introduction

Throughout adolescence, individuals undergo extensive maturation, marked by major neurodevelopment and behavioral changes as they transition into adulthood. During this period, adolescents show increased risk-taking and reward-seeking behaviors, especially the initiation of drug use (Casey, Getz, & Galvan, 2008; Masten, Faden, Zucker, & Spear, 2009; Spear, 2000). Alcohol, in particular, is consumed at alarming rates during adolescence, with 7.4 million individuals under the age of twenty-one reporting drinking in the past month (SAMHSA, 2019). Young adolescence in particular appears to be a crucial time for the initiation of problematic alcohol use; many individuals who begin drinking early go on to rapidly escalate their use, with more than ten percent accelerating to meet alcohol use disorder (AUD) criteria within the first year of initiation (Faden, 2006; Forman-Hoffman, Edlund, Glasheen, & Ridenour, 2017; Masten et al., 2009). This early introduction to alcohol use has been associated with numerous long-lasting alterations to brain function, structure, and behavior (Crews, Vetreno, Broadwater, & Robinson, 2016; Spear, 2018; Squeglia, Jacobus, & Tapert, 2014), including increased rates of anxiety, depression, and AUD later on (Duncan, Alpert, Duncan, & Hops, 1997; Grant & Dawson, 1997; Hawkins et al., 1997; Pitkanen, Lyyra, & Pulkkinen, 2005; Sung, Erkanli, Angold, & Costello, 2004; Wolford-Clevenger & Cropsey, 2019). Our understanding of the effects of adolescent alcohol comes largely from studies that have used exposures yielding BECs of 150-250 mg/dL, far exceeding the definition of binge-level exposure [80 mg/dL, (NIAAA, 2004)]. However, adolescents do not routinely drink to these levels (SAMHSA, 2019). Thus, there is a need to further examine the impacts of moderate levels of intoxication, as these are understudied and still relevant to this population.

Animal models have been used to examine the long-term effects of adolescent alcohol use and have found interesting, yet somewhat inconclusive, results regarding negative affect and reward-related behaviors. Some research has shown increased adult voluntary drinking and anxiety-like behavior following adolescent ethanol exposure (Alaux-Cantin et al., 2013; Kyzar, Zhang, & Pandey, 2019; Pandey, Sakharkar, Tang, & Zhang, 2015; Sakharkar et al., 2019), whereas others have identified decreases or no changes in these behaviors (Gass et al., 2014; Nentwig, Starr, Chandler, & Glover, 2019; Torcaso, Asimes, Meagher, & Pak, 2017). Of the few preclinical studies that have looked at the effects of low to moderate levels of intoxication, some results appear to be relatively consistent with the higher-level exposures, showing increased voluntary drinking in adulthood (Amodeo, Kneiber, Wills, & Ehlers, 2017; Broadwater, Varlinskaya, & Spear, 2013). However, lower levels of exposure have failed to identify shortterm changes in anxiety-like behavior in adolescent, adult, or aged rats (Matthews et al., 2019), but it is not clear how these results may translate to long-term effects. Taken together, adolescent ethanol exposure appears to have long-term consequences on several affective and reward-related behaviors, but the ways in which this exposure impacts the underlying neural mechanisms responsible for changes in these behaviors is still not well understood, especially with moderate-level exposures.

One brain region that plays a major role in affective disruption and reward processing is the basolateral nucleus of the amygdala (BLA). This region receives input from sensory and cortical areas and sends direct projections to the extended amygdala, the major amygdalar output regions, for responses to emotional stimuli (Janak & Tye, 2015; Tye et al., 2011). The BLA is involved in the modulation of many negative affective behaviors, including anxiety, namely through the balance of excitatory glutamatergic and inhibitory GABAergic neurotransmission in the region (Aroniadou-Anderjaska, Qashu, & Braga, 2007; Janak & Tye, 2015; Shekhar, Truitt, Rainnie, & Sajdyk, 2005; Tye et al., 2011). Disruption of this balance, particularly decreased inhibitory or increased excitatory modulation of the BLA, leads to overexcitation and ultimately increased anxiety-like responses (Sajdyk & Shekhar, 1997, 2000; Shekhar et al., 2005). In studies that have examined changes to synaptic activity in the BLA during acute withdrawal from ethanol exposure during adolescence (based on weight), increased glutamate transmission (Christian, Alexander, Diaz, Robinson, & McCool, 2012; Läck, Diaz, Chappell, DuBois, & McCool, 2007; McGinnis, Parrish, Chappell, Alexander, & McCool, 2020; McGinnis, Parrish, & McCool, 2020; Morales, McGinnis, Robinson, Chappell, & McCool, 2018) and decreased GABA transmission (Diaz, Christian, Anderson, & McCool, 2011) were found. This indicates that in the early stages of withdrawal following adolescent ethanol exposure, the excitatory/inhibitory balance is shifted within the BLA, which ultimately produces anxiety-like behavior. Whether this shift in synaptic function is maintained into adulthood has yet to be examined. Interestingly, adolescent ethanol exposure produces alterations to histone modification and DNA methylation mechanisms in the central amygdala, but not the BLA, which have been linked to increased anxiety-like behaviors (Pandey et al., 2015; Sakharkar et al., 2019). However, functional changes in the BLA caused by adolescent ethanol exposure have been implicated in affective and reward-related dysfunction in adulthood (Moaddab, Mangone, Ray, & McDannald, 2017). This suggests that mechanisms in the BLA independent of epigenetic processes, such as synaptic transmission, may be disrupted long term.

While the glutamate and GABA systems directly shift the excitatory/inhibitory balance within the BLA, there are several neuromodulators that fine-tune this balance and which are targets of alcohol. One such system that is highly expressed in the BLA is the kappa opioid receptor (KOR) system. This system has long been implicated in negative affective states (Hang, Wang, He, & Liu, 2015; Van’t Veer & Carlezon, 2013) and modulation of reward systems within the brain (Bruijnzeel, 2009). This system is also a target of ethanol, as release of dynorphin, the endogenous ligand for the KOR, occurs in several brain regions following acute ethanol administration (Jarjour, Bai, & Gianoulakis, 2009; Lam, Marinelli, Bai, & Gianoulakis, 2008; Marinelli, Lam, Bai, Quirion, & Gianoulakis, 2006). Chronic exposure to ethanol also targets this system, such that adaptations to the KOR system have been identified following this exposure. A general upregulation of KOR/dynorphin mRNA and protein has been demonstrated across the brain following exposure to ethanol in adulthood [for review, see (Anderson & Becker, 2017; Diaz, Przybysz, & Rouzer, 2017)]. In the extended amygdala, functional adaptations also occur following chronic exposure. For example, chronic ethanol exposure increases the sensitivity of KORs within the nucleus accumbens, which plays a direct role in regulation of dopamine release (Karkhanis, Huggins, Rose, & Jones, 2016; Rose et al., 2016). In the central amygdala, increased levels of dynorphin release and receptor coupling have been linked to elevated alcohol consumption and negative affect following chronic alcohol exposure (Kissler et al., 2014; Kissler & Walker, 2016). Finally, blockade of KORs in the BNST reduced alcohol consumption, negative affect, and physiological withdrawal symptoms following chronic alcohol exposure (Erikson, Wei, & Walker, 2018), while activation of KORs in this region enhanced alcohol seeking following chronic alcohol exposure (Funk, Coen, Tamadon, & Le, 2019). Together, these studies clearly demonstrate that parts of the extended amygdala which regulate negative affective states and modulation of reward have disrupted KOR function following chronic alcohol exposure; however, whether similar adaptations occur in the BLA, particularly in response to exposure in adolescence, is unknown.

The similarities in behavioral alterations following chronic alcohol exposure in both adolescents and adults and the adaptations in the KOR system in response to adult alcohol exposure suggest that KOR function is likely also altered by chronic exposure to alcohol in adolescence. Importantly, we have shown that KOR function is developmentally regulated within the BLA (Przybysz, Werner, & Diaz, 2017) and mounting evidence suggests that alcohol exposure may differentially affect the KOR system depending on the developmental period in which exposure occurs [for review, see: (Diaz et al., 2017)]. Furthermore, several studies have also demonstrated that adolescent alcohol exposure can disrupt developmental processes that occur between adolescence and adulthood (Spear & Swartzwelder, 2014). Therefore, we sought to determine whether exposure to moderate levels of alcohol during adolescence would alter adult KOR function and lead to further disruptions in the excitatory/inhibitory balance within the BLA.

## Methods

### Animals

This study included male and female Sprague-Dawley rats that were bred and reared in our animal colony at Binghamton University, with breeding pairs originating from Envigo (Indianapolis, IN). Animals were housed in a temperature-controlled (22□C) vivarium, given *ad libitum* access to standard lab chow (LabDiet 5L0D. PicoLab Laboratory Rodent Diet, ScottPharma Solutions, Marlborough, MA) and water, and maintained on a 12:12 h light: dark cycle (lights on at 0700 h). Litters were culled to 12 pups (six males and six females) whenever possible on postnatal day (P) 2 and reared with their mothers until weaning on P21. Following weaning, animals were group-housed in groups of 2-3 animals of each sex per cage. All animal procedures were approved by the Binghamton University Institutional Animal Care and Use Committee.

### Moderate adolescent Chronic Intermittent Ethanol (aCIE) Exposure

Starting on P30, animals in their home cages were transferred to vapor inhalation chambers for ten consecutive days. Animals were exposed to either room air (control group) or vaporized ethanol (aCIE group) for 12 h overnight (2000-0800 h) every other night during this ten-day period, resulting in five cycles of 12 h ethanol on, 36 h ethanol off. All animal husbandry (food, water, and bedding changes) occurred during the “off” periods, and no handling or movement of the animals occurred during the 12-hour exposure periods. Vapor levels were recorded just prior to the end of each exposure day to ensure consistent exposure throughout the 10-day exposure period and across cohorts. Rat weight was also monitored throughout the exposure period, with rats being weighed before being placed into the vapor chambers, twice at regular intervals during the 10-day exposure period, and following the end of the last exposure. Rat chow was replaced after each exposure to ethanol to avoid consumption of ethanol saturated chow during the periods of abstinence. Following the final cycle, animals were transferred back to the animal colony until adulthood (~P70).

### BEC Analysis

Blood ethanol concentrations (BECs) were determined in a separate subset of animals. This set of animals was exposed on a reverse cycle (0800 h to 2000 h) to collect blood samples during the 12-hour exposure period, with tail blood samples taken every two hours on day 1 and day 9 of the exposure. All animals used for BEC determination were euthanized following the day of blood collection. BECs were determined using an Analox AM-1 alcohol analyzer (Analox Instruments, Lunenburg, MA). The machine was calibrated to a 100 mg% industry standard and was re-calibrated every ten samples. All samples were run in duplicate and an average reading was recorded for each pair in mg/dL. The floor of assay sensitivity is ~15 mg/dL as evidenced by samples collected prior to alcohol exposure, thus all measurements below this threshold were interpreted as values of zero.

### Whole-cell patch-clamp electrophysiology

All electrophysiological procedures were done as previously described (Przybysz et al., 2017). Briefly, adult rats (P70-100) were sedated with ketamine (250 mg/kg) and quickly decapitated. Brains were rapidly removed and immersed in cold oxygenated (95% 02, 5% CO2) sucrose artificial cerebrospinal fluid (ACSF) cutting solution containing (in mM): sucrose (220), KCl (2), NaH2PO4 (1.25), NaHCO3 (26), glucose (10), MgSO4 (12), CaCl2 (0.2), and ketamine (0.43). 300 μm brain slices containing BLA were made using a Vibratome (Leica Microsystems, Bannockburn, IL, USA). Slices were incubated in oxygenated normal ACSF containing (in mM): NaCl (126), KCl (2), NaH2PO4 (1.25), NaHCO3 (26), glucose (10), CaCl2 (2), MgSO4 (1), and ascorbic acid (0.4), and allowed to recover for at least 40 min at 34°C before recording. Slices remained at 34°C and all experiments were performed 1-4 h after slice preparation.

Following incubation, slices were transferred to a recording chamber in which oxygenated ACSF was warmed to 32°C and superfused over the slice at 3 mL/min. All recordings were made from pyramidal neurons within the BLA that were visualized using infrared-differential interference contrast microscopy (Olympus America, Center Valley, PA). Pyramidal neurons were identified based on morphology and capacitance (> 150 pF) as previously described (Przybysz et al., 2017). Spontaneous inhibitory postsynaptic current (sIPSC) recordings were collected with patch pipettes filled with KCl internal solution containing (in mM): KCl (135), HEPES (10), MgCl2 (2), EGTA (0.5), Mg-ATP (5), Na-GTP (1), and QX314-Cl (1), pH of 7.25, and osmolarity of 280-290 mOsm. Spontaneous excitatory postsynaptic current (sEPSC) recordings were made with a K-gluconate internal solution, containing (in mM): K-Gluconate (120), KCl (15), EGTA (0.1), HEPES (10), MgCl2 (4), MgATP (4), Na3GTP (0.3), phosphocreatine (7), and QX-314 Br (1.5), pH of 7.3, and osmolarity of 295-305 mOsm. Data were acquired with a MultiClamp 700B (Molecular Devices, Sunnyvale, CA) at 10 kHz, filtered at 1 kHz, and stored for later analysis using pClamp software (Molecular Devices). For recordings of GABA_A_ receptor-mediated sIPSCs, we pharmacologically blocked AMPA and NMDA glutamate receptors using 1 mM kynurenic acid and 50 μM DL-APV, respectively. For sEPSC recordings, we blocked GABA_A_ receptors with 10 mM gabazine.

Upon opening, neurons were allowed to equilibrate for at least 5 min before a baseline was recorded. We have previously shown that the KOR agonist U69593 is effective ~1 min into drug application (Przybysz et al., 2017; Varlinskaya, Johnson, Przybysz, Deak, & Diaz, 2020), so following 3 min of baseline recording, we recorded at least 4 min of continuous U69593 (1 μM) application. Given potential variability in timing of U69593 reaching the slice and instability of the baseline in the first 1 minute of recording, we discarded the first minute of recording and only used the subsequent minute to quantify the baseline (see time-courses below). Access resistance was monitored throughout each recording, and only recordings in which the access resistance changed <20% were kept for analysis.

### Drugs and Chemicals

All chemicals were purchased from Sigma-Aldrich (St. Louis, MO) unless otherwise noted. Kynurenic acid, DL-APV, and QX314-Cl were purchased from Tocris/R&D Systems (Bristol, UK).

### Statistics

Data were analyzed using MiniAnalysis (Synaptosoft Inc.) and were analyzed statistically with Prism 8 (GraphPad, San Diego, CA). Given evidence that BLA physiology is substantially different between males and females (Blume et al., 2017), data from males and females were analyzed separately. Sample sizes indicate number of individual cells, but no more than 2 cells were recorded from a single animal in any given experiment. Data were first analyzed using the Pearson omnibus and K-S normality tests. If data followed a normal distribution, parametric tests were used; otherwise, nonparametric tests were used. Grubbs outlier tests were run on all basal and U69593 data sets, and any identified outliers were removed from all analyses. To assess changes in BECs across exposure on the first and last days of exposure in males and females, two-way repeated-measures ANOVAs (day of exposure x hour of exposure) were used followed by Fisher’s LSD post hoc test. For U69593 experiments, K-S tests to compare distributions were used to determine the responsiveness of individual cells to the drug. All data are presented as mean ± SEM, with p < 0.05 considered statistically significant.

## Results

### aCIE Exposure Characterization

Blood samples were collected at two-hour intervals, beginning at 08:00 and ending at 20:00, on the first and last day of the exposure period. For both males and females, BECs on day 1 rose continuously. For males **(Fig. 1A)**, this exposure yielded BECs that peaked at the 12^th^ hour of exposure with an average of 109.8 ± 11.71 mg/dL at this timepoint. On the last day of exposure, male BECs also peaked at the 12^th^ hour (59.94 ± 4.00 mg/dL), but there was a significant decrease in BECs between the first and final days of exposure (*F*(1,14) = 34.54, *n* = 16, *p* < 0.0001); follow-up with Fisher’s LSD post-hoc test revealed significantly lower average BECs on the final exposure day for hours: four (*p* = 0.035), eight (*p* = 0.001), ten (*p* = 0.002), and twelve (*p* = 0.003). For females **(Fig. 1B)**, the first day of exposure yielded BECs that also peaked at hour 12 with an average of 96.17 ± 17.16 mg/dL at this timepoint. Females also exhibited a decrease in BECs between the first and last days of exposure, with female BECs averaging 50.21 ± 7.18 mg/dL at the 12^th^ hour of the final exposure. This also indicated a significant decrease in BECs between first and final exposure (*F*(1,14) = 5.030, *n* = 16, *p* = 0.042); followup with Fisher’s LSD post-hoc revealed significantly lower BECs during the final exposure day for hours four (*p* = 0.003), six (*p* < 0.0001), eight (*p* = 0.002), ten (*p* < 0.0001), and twelve (*p* < 0.0001). Male and female rat weights were taken before, during, and after the exposure period in both air and aCIE exposed groups. For males **(Fig. 1C)**, there was a significant main effect of day (*F*(1.100, 60.89) = 1706, *N* = 66, *p* < 0.0001). There was no main effect of exposure (*F*(1, 64) = 0.22, *N* = 66, *p* = 0.64). There was a significant day x exposure interaction (*F*(3,166) = 6.45, *N* = 66, *p* = 0.004), but Fisher’s LSD post-hoc showed no significant differences in weight between exposure groups at any individual exposure day. For females **(Fig. 1D)**, there was a significant main effect of day (*F*(1.629, 91.77) = 1056, *N* = 65, *p* < 0.0001), a significant main effect of exposure (*F*(1.63) = 5.82, *N* = 65, *p* = 0.02), and a significant day x exposure interaction (*F*(3, 169) = 8.6, *N* = 65, *p* < 0.0001). Post-hoc tests revealed that aCIE females weighed significantly less than air females on P33 (*p* = 0.018) and P37 (*p* = 0.019), but these differences were no longer present by P40 (*p* = 0.419). Ethanol vapor levels were recorded just prior to the end of each 12-hour ethanol exposure, and analysis of these indicated that vapor levels were consistent across all exposure days and cohorts of animals (mean vapor level: 4.2 g/L ± 0.134, data not shown).

**Figure 1.**
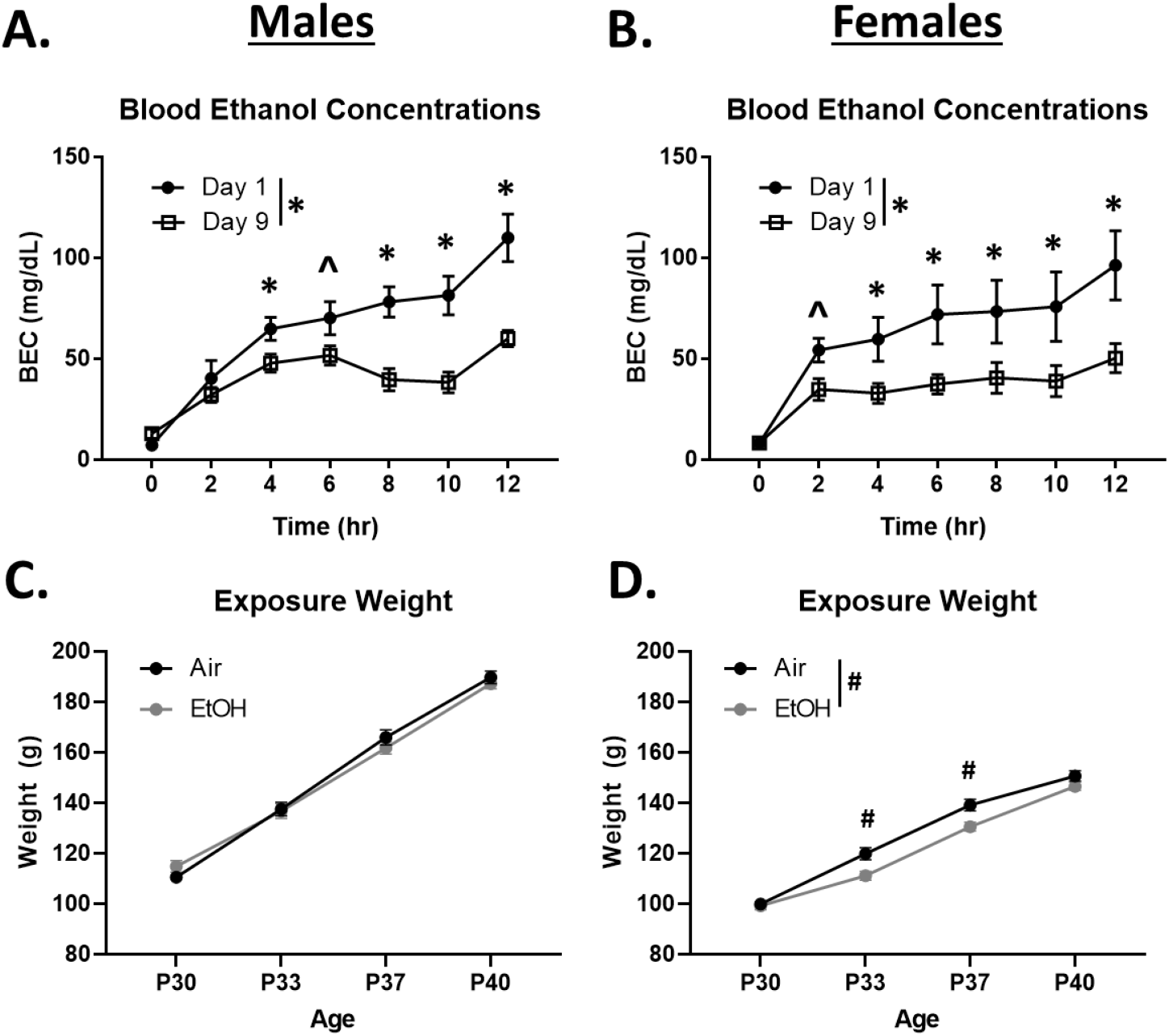
Characterization of aCIE exposure paradigm. (A, B) Blood ethanol concentration curves across the 12-hour exposure on the first day (day 1) and last day (day 9) of exposure for males (left) and females (right). * = p < 0.05, ^ = 0 < 0.08 between day 1 and 9. (C, D) Change in weight across exposure days in air and aCIE-exposed males (left) and females (right). # = p < 0.05 between air and aCIE groups.

### Basal GABA transmission in adult BLA

We performed electrophysiological experiments in the BLA of adult rats that had undergone the aCIE exposure paradigm. Assessment of basal GABA transmission in the BLA of males showed that aCIE exposure did not alter sIPSC frequency (**Fig. 2A**: *t* = 1.69, *df* = 14, *n* = 7 – 9, *p* = 0.11) or amplitude (**Fig. 2B**: *t* = 0.11, *df* = 14, *n* = 7 – 9, *p* = 0.91). In females, there was also no effect of aCIE on sIPSC frequency (**Fig. 2C**: *t* = 0.15, *df* = 13, *n* = 7 – 8, *p* = 0.88) or amplitude (**Fig. 2D**: *t* = 0.21, *df* = 13, *n* = 7 – 8, *p* = 0.84).

**Figure 2.**
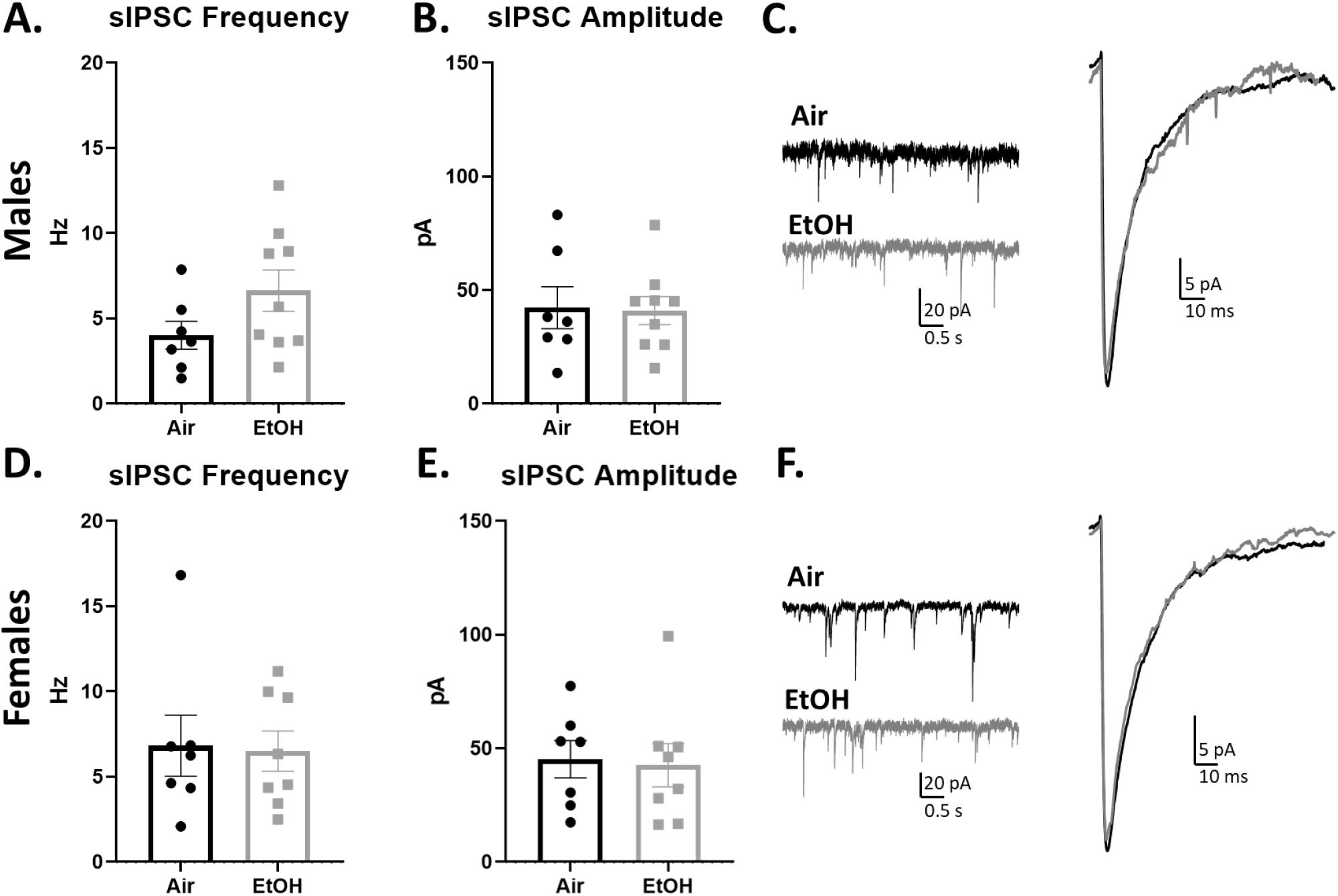
Exposure to aCIE does not alter adult BLA GABA transmission. Comparison of BLA sIPSC frequency (A, D) and amplitude (B, E) between air and aCIE-exposed adult males (top) and females (bottom), with exemplar traces demonstrating no effect of aCIE exposure in either sex (C, F). Bars depict mean ± SEM.

### aCIE shifts GABA responsiveness to U69593

We next assessed the effect of U69593 (1 μM) on GABA transmission within the BLA. When averaging across all cells, in neurons from air-exposed males there was no effect of 1 μM U69593 on sIPSC frequency (**Fig. 3A**: 6.81 ± 15.77% change from baseline, *n* = 7), consistent with our previously reported findings from naïve adult males using 1 μM U69593 (Przybysz et al., 2017). However, in ethanol-exposed males, U69593 significantly decreased sIPSC frequency (**Fig 3A**: −17.23 ± 6.48% change from baseline, *n* = 9, *p* = 0.03 compared to 0). There was no effect of U69593 on sIPSC amplitude in air- or ethanol-exposed males (**Fig. 3B**: *air*: 24.41 ± 15.28% change from baseline, *n* = 7, *p* = 0.22 compared to 0; *EtOH:* −3.47 ± 7.05% change from baseline, *n* = 9, *p* = 0.73 compared to 0). Despite these aCIE-induced differences in U69593 modulation of sIPSC frequency, we observed high variability in controls, consistent with our previous observations (Przybysz et al., 2017). Therefore, to determine whether aCIE exposure shifted the responsiveness of individual cells to U69593, we ran Kolmogorov-Smirnov (K-S) tests on each recording as we have done previously (Varlinskaya et al., 2020). This analysis revealed that the aCIE exposure shifted the proportion of cells that responded to U69593. In air-exposed males, 3/7 cells were non-responsive, while sIPSC frequency was inhibited in 2/7 cells and was potentiated in 2/7 cells. However, in EtOH-exposed males, 4/9 cells were nonresponsive, but sIPSC frequency was inhibited in all other cells (5/9); no cells were potentiated by U69593. Bar graphs illustrating this shift are shown in **Fig. 3C**, and time courses for each of the groups from air- and EtOH-exposed animals are displayed in **Fig. 3D** and **3E**, respectively.

**Figure 3.**
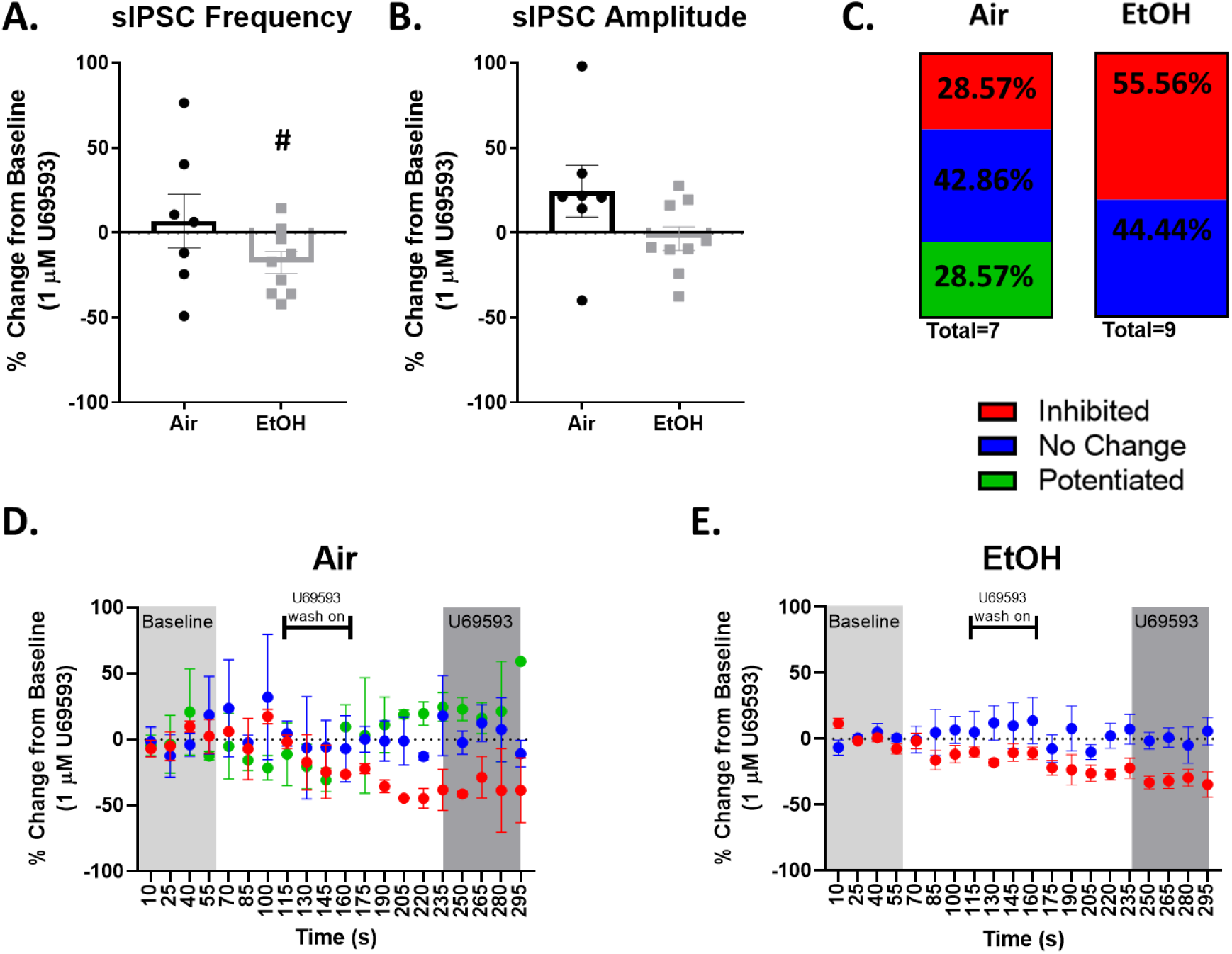
Exposure to aCIE alters KOR modulation of GABA transmission and individual cell responsiveness to KOR activation in adult male BLA. Change from baseline (%) of sIPSC frequency (A) and amplitude (B) for all cells following application of KOR agonist U69593 (1 μM) in air- and aCIE-exposed males. # = p < 0.05, significant change from baseline. (C) Component chart indicating an aCIE-induced shift in the percentage of BLA pyramidal cells that were inhibited (red), not changed (blue), or potentiated (green) by KOR activation. (D, E) Frequency time-courses of U69593 application for air-(D) or aCIE-(E) exposed males separated by directional change. Light gray and dark gray shaded boxes correspond to the 60 s period used to calculate baseline and % change from baseline, respectively, in A, and bracket indicates the period when U69593 began washing onto the slice.

In neurons from females, U69593 had no effect on sIPSC frequency (**Fig. 4A**: *air:* −6.85 ± 8.81% change from baseline, *n* = 8, *p* = 0.46 compared to 0; *EtOH:* 0.54 ± 6.35% change from baseline, *n* = 8, *p* = 0.93 compared to 0) or amplitude (**Fig 4B:** *air:* −6.89 ± 13.26% change from baseline, *n* = 8, *p* = 0.62 compared to 0; *EtOH:* −2.62 ± 11.16% change from baseline, *n* = 8, *p* = 0.92 compared to 0). However, when analyzing for shifts in individual cell responsivity using the K-S test, we once again found evidence for an aCIE-induced shift. In air-exposed females, the majority of cells were inhibited by U69593 (5/8), while one cell was potentiated by U69593 and 2 cells were nonresponsive. In EtOH-exposed females, this proportion was shifted so that the majority of cells were nonresponsive (6/8), one cell was inhibited, and one cell was potentiated. Bar graphs and time courses illustrating this shift can be found in **Fig. 4C**, **4D,** and **4E**.

**Figure 4.**
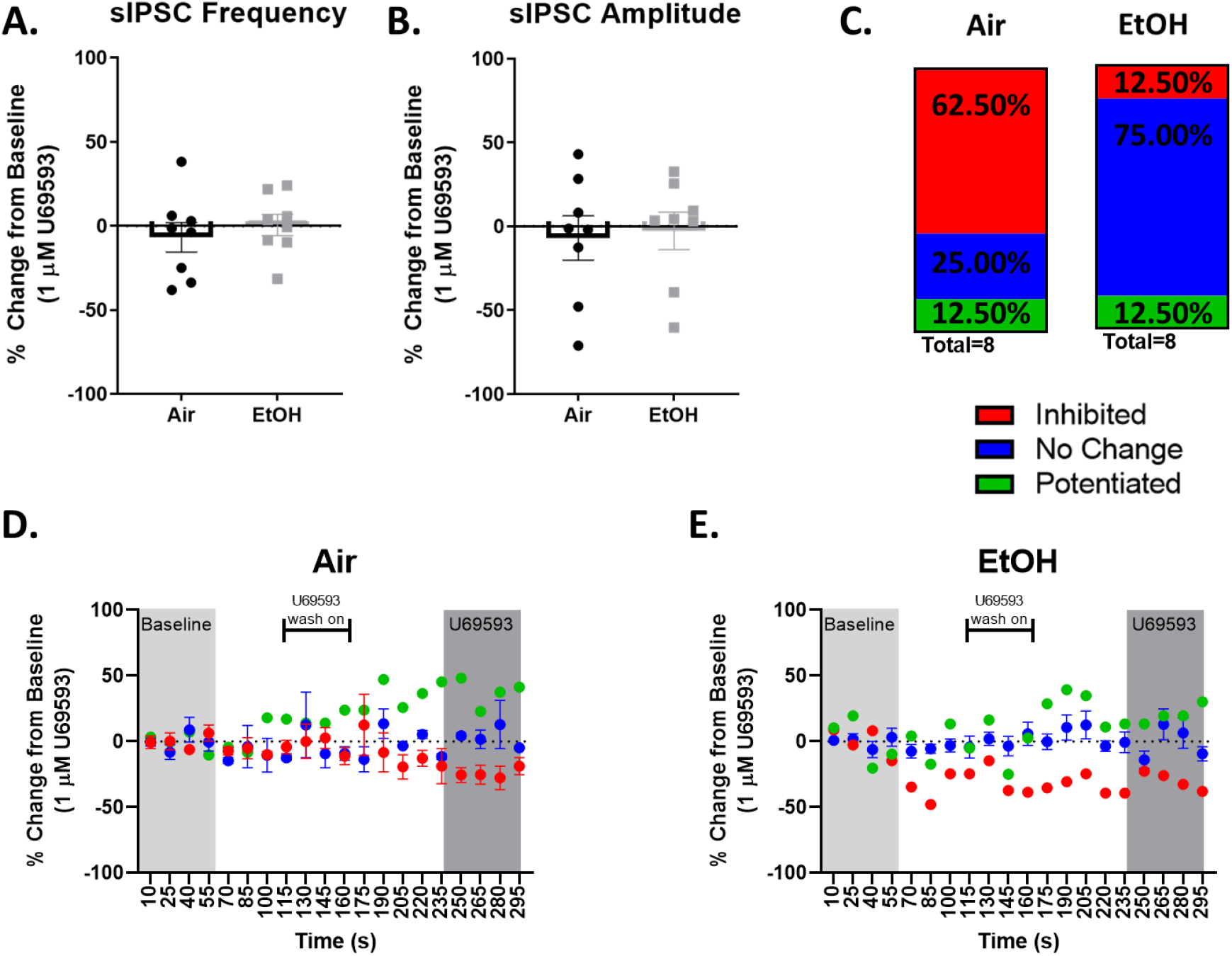
Exposure to aCIE does not alter KOR modulation of GABA transmission but shifts individual cell responsiveness to KOR activation in adult female BLA. Change from baseline (%) of sIPSC frequency (A) and amplitude (B) for all cells following application of KOR agonist U69593 (1 μM) in air- and aCIE-exposed females. C) Component chart indicating an aCIE-induced shift in the percentage of BLA pyramidal cells that were inhibited (red), not changed (blue), or potentiated (green) by KOR activation. (D, E) Frequency time-courses of U69593 application for air-(D) or aCIE-(E) exposed females separated by directional change. Light gray and dark gray shaded boxes correspond to the 60 s period used to calculate baseline and % change from baseline, respectively, in A, and bracket indicates the period when U69593 began washing onto the slice.

### Basal glutamate transmission in adult BLA

Assessment of aCIE exposure effects on glutamate transmission in the BLA showed that in males neither sEPSC frequency (**Fig. 5A**: *t* = 0.78, *df* = 9, *n* = 5 – 6, *p* = 0.45) or amplitude (**Fig. 5B**: *t* = 0.35, *df* = 9, *n* = 5 – 6, *p* = 0.74) was not affected by aCIE. However, in females, aCIE exposure significantly increased sEPSC frequency (**Fig. 5C**: *t* = 2.22, *df* = 14, *n* = 8 – 9, *p* = 0.04), with a trend toward increasing sEPSC amplitude (**Fig. 5D**: *t* = 1.92, *df* = 14, *n* = 8 – 9, *p* = 0.07).

**Figure 5.**
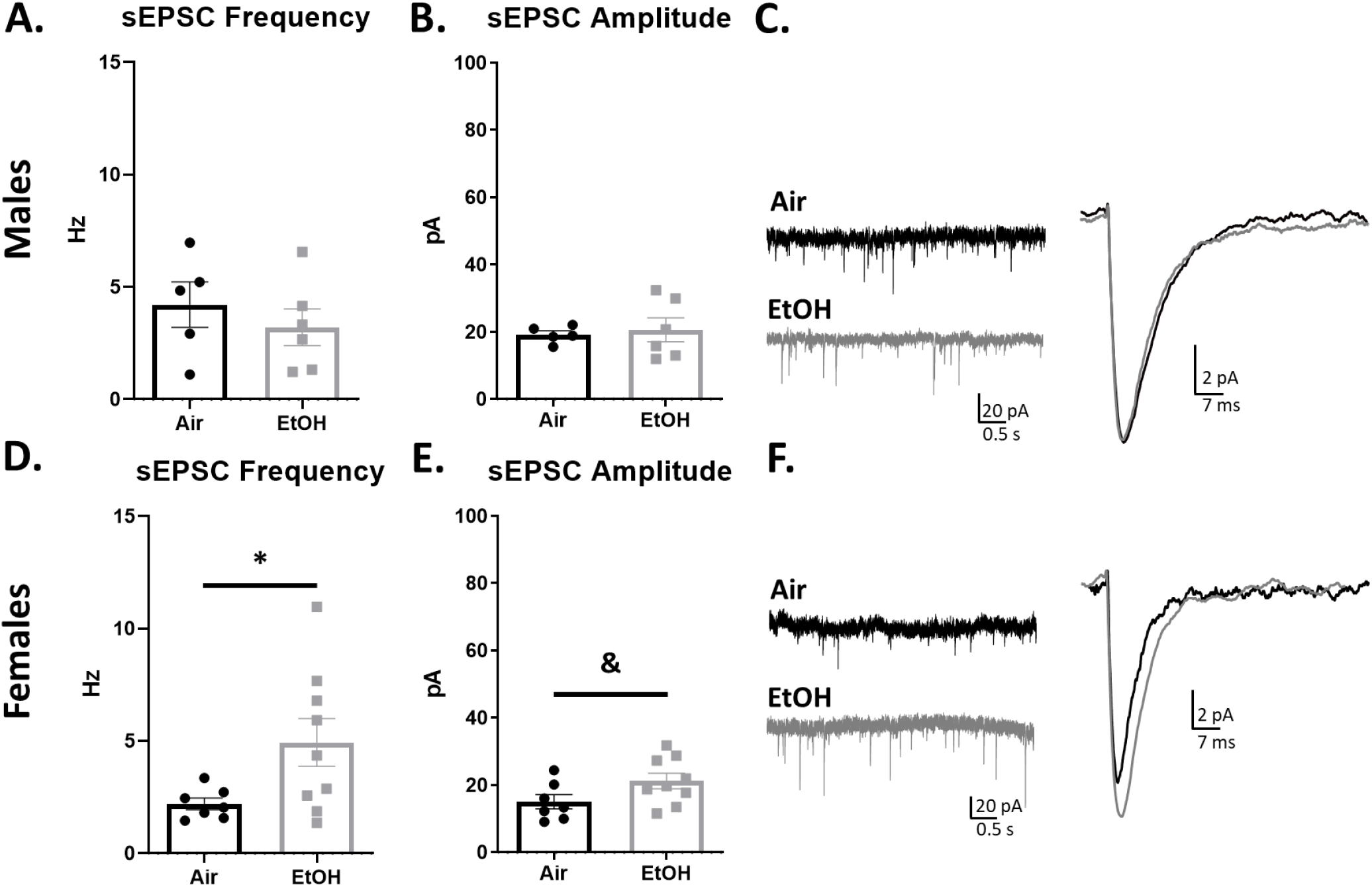
Exposure to aCIE increases adult female BLA glutamate transmission. Comparison of BLA sEPSC frequency (A, D) and amplitude (B, E) between air- and aCIE-exposed adult males (top) and females (bottom), * = p < 0.05 between air and aCIE groups, & = p = 0.07 between air and aCIE groups. (C, F) Exemplar traces demonstrating no aCIE-induced difference in males (C) but a significant increase in sEPSC frequency and a trend toward increased sEPSC amplitude in aCIE females (F).

### U69593-mediated glutamate transmission

When assessing the effect of U69593 on BLA glutamate transmission, we found no effect on sEPSC frequency in air- or ethanol-exposed males (**Fig. 6A**: *air:* −4.18 ± 10.963% change from baseline, *n* = 5, *p* = 0.72 compared to 0; *EtOH:* 10.94 ± 15.90 % change from baseline, *n* = 6, *p* = 0.52 compared to 0). However, while U69593 did not affect sEPSC amplitude in air-exposed males (**Fig. 6B**: −18.73 ± 6.97 % change from baseline, *n* = 5, *p* = 0.13 compared to 0), it did significantly decrease sEPSC amplitude in EtOH-exposed males (**Fig 6B**: −8.77 ± 5.32 % change from baseline, *n* = 6, *p* = 0.03 compared to 0). When using the K-S test to assess shift in responsivity to U69593, we found a slight change caused by aCIE exposure. In air-exposed males, all cells but one were nonresponsive (4/5), with the one responsive cell being inhibited.

**Figure 6.**
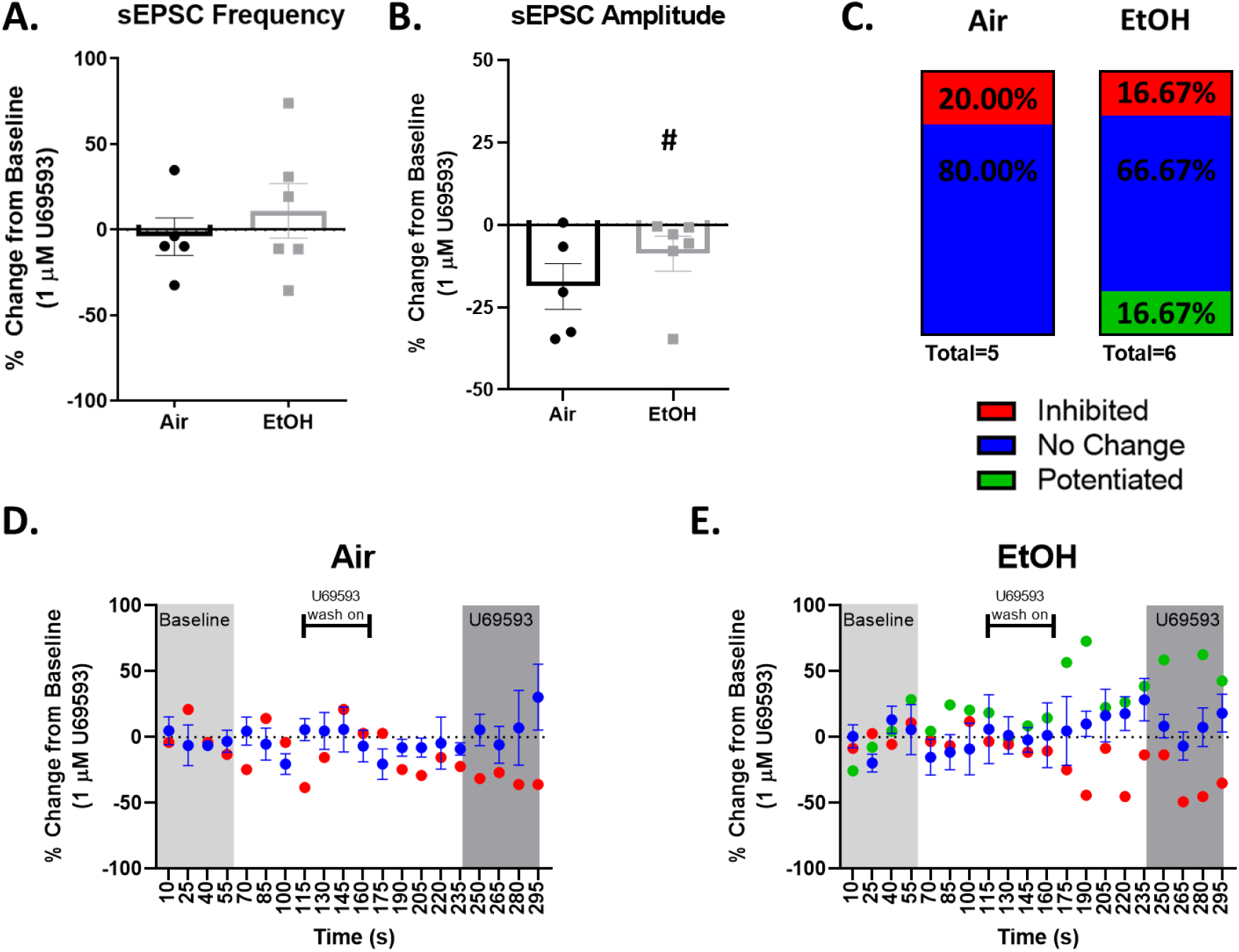
KOR activation inhibits BLA glutamate transmission in aCIE-exposed adult males. Change from baseline (%) of sEPSC frequency (A) and amplitude (B) for all cells following application of KOR agonist U69593 (1 μM) in air- and aCIE-exposed males. # = p < 0.05, significant change from baseline. (C) Component chart indicating an aCIE-induced shift in the percentage of BLA pyramidal cells that were inhibited (red), not changed (blue), or potentiated (green) by KOR activation. (D, E) Frequency time-courses of U69593 application for air-(D) or aCIE-(E) exposed males separated by directional change. Light gray and dark gray shaded boxes correspond to the 60 s period used to calculate baseline and % change from baseline, respectively, in A, and bracket indicates the period when U69593 began washing onto the slice.

In EtOH-exposed males, all but two cells were nonresponsive (4/6), with one cell being inhibited and one cell being potentiated.

In both air- and ethanol-exposed females, U69593 had no effect on sEPSC frequency (**Fig. 7A**: *air*. −13.56 ± 9.33% change from baseline, *n* = 8, *p* = 0.20 compared to 0; *EtOH:* −5.94 ± 9.46% change from baseline, *n* = 9, *p* = 0.55 compared to 0) or sEPSC amplitude (**Fig. 7B**: *air*: −2.65 ± 5.47% change from baseline, *n* = 8, *p* = 0.65 compared to 0; *EtOH:* −7.49 ± 3.76% change from baseline, *n* = 9, *p* = 0.09 compared to 0). When analyzing individual cell responsiveness to U69593 using the K-S test, we once again found a slight aCIE exposure-induced shift. The majority of cells from air-exposed females were nonresponsive (5/7), with the other two cells both being inhibited. In ethanol-exposed females, 4/8 cells were nonresponsive, while 2/8 cells were inhibited by U69593 and 2/8 cells were potentiated by U69593.

**Figure 7.**
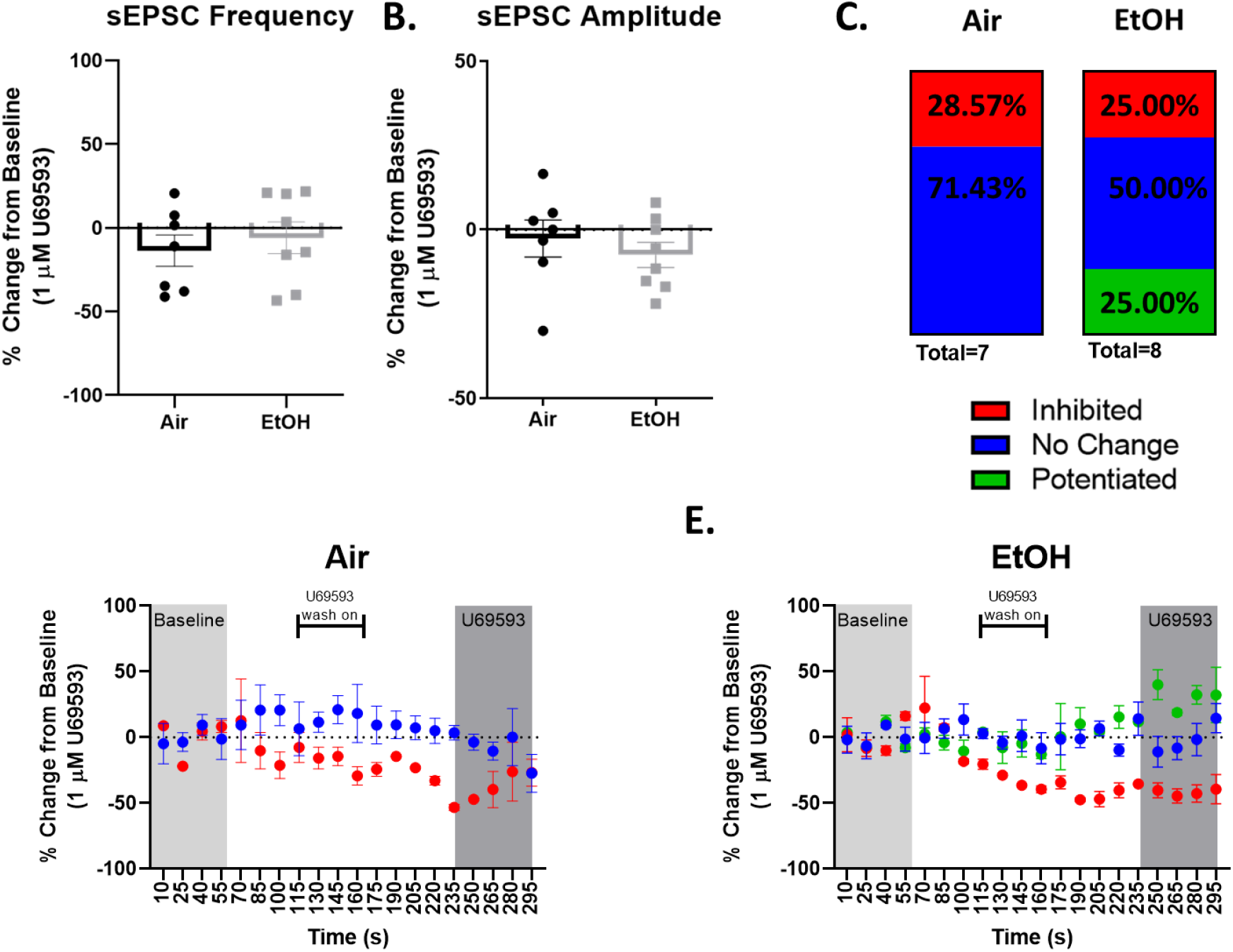
Exposure to aCIE does not alter KOR modulation of glutamate transmission but shifts individual cell responsiveness to KOR activation in adult female BLA. Change from baseline (%) of sEPSC frequency (A) and amplitude (B) for all cells following application of KOR agonist U69593 (1 μM) in air- and aCIE-exposed females. (C Component chart indicating an aCIE-induced shift in the percentage of BLA pyramidal cells that were inhibited (red), not changed (blue), or potentiated (green) by KOR activation. (D, E) Frequency time-courses of U69593 application for air-(D) or aCIE-(E) exposed females separated by directional change. Light gray and dark gray shaded boxes correspond to the 60 s period used to calculate baseline and % change from baseline, respectively, in A, and bracket indicates the period when U69593 began washing onto the slice.

## Discussion

Numerous studies have demonstrated that exposure to alcohol during adolescence has long-term consequences, including increased risk for developing an anxiety disorder or alcohol use disorder in adulthood (Crews et al., 2016). Despite the well-established role of the BLA in disorders of this type, few studies have investigated the long-term effects of adolescent alcohol exposure on BLA physiology, particularly using moderate levels of ethanol. Therefore, one goal of the present study was to understand how exposure to alcohol during adolescence may affect adult inhibitory (GABAergic) and excitatory (glutamatergic) neurotransmission in the BLA. Our findings indicate that adult males who had gone through aCIE exposure exhibited no changes to basal glutamate transmission or basal GABA transmission compared to air-exposed controls. Females exposed to aCIE also showed no change in GABA transmission but did exhibit a significant increase in sEPSC frequency and a trend toward an increase in sEPSC amplitude compared to air-exposed controls, together suggesting a persistent increase in glutamate transmission in females following adolescent alcohol exposure.

The present work is the first to examine long-term changes to BLA physiology following alcohol exposure during adolescence. However, studies examining the effects of adolescent vapor exposure followed by acute withdrawal have provided evidence that adolescent alcohol exposure disrupts excitatory and inhibitory transmission in the male BLA. For example, one study that exposed adolescent (based on weight) male Sprague Dawley rats to intermittent vapor ethanol for 10 days found enhanced presynaptic glutamate release and enhanced postsynaptic NMDA receptor function 24 hours into withdrawal (Läck et al., 2007). This group has also identified different alterations on specific glutamatergic inputs into the BLA, finding that projections from the prefrontal cortex (Christian et al., 2012) and agranular insular cortex (McGinnis, Parrish, & McCool, 2020) have enhanced postsynaptic AMPA receptor function. They also found that glutamatergic projections from the dorsal and ventral medial prefrontal cortex (mPFC) to the BLA are differentially impacted, showing that presynaptic glutamate release from the dorsal mPFC is increased while it is decreased from the ventral mPFC 24 hours into withdrawal (McGinnis, Parrish, Chappell, et al., 2020). While our sEPSC findings in males do not reflect those of these studies, a major difference is the period of abstinence from ethanol, as our rats underwent prolonged (30+ days) abstinence following exposure. Two studies that exposed adolescent rats to eight cycles of 16 hours on and 8 hours off from vaporized ethanol found no apparent effects on NMDA receptor-mediated long term depression in the BNST 30 days after exposure (Carzoli et al., 2019; Kasten et al., 2020), despite observing alterations to this and increased sEPSC frequency 24 hours after exposure (Carzoli et al., 2019). Based on these studies in the BNST and our own findings, it seems likely that the disrupted mechanisms identified during acute withdrawal undergo additional compensatory adaptations during protracted abstinence. In the present study, we did not examine any potential changes to intrinsic excitability of BLA neurons, nor did we explore possible input-specific or population-specific effects as was done in those studies described above. It is possible that alterations may depend on the neuronal population or specific synapses being sampled, as these previous studies directly examined (Christian, Alexander, Diaz, & McCool, 2013; Christian et al., 2012; McGinnis, Parrish, Chappell, et al., 2020; McGinnis, Parrish, & McCool, 2020; Morales et al., 2018). Additionally, the studies outlined above used a greater total number of exposures or had exposures that induced greater BECs in their animals compared to our aCIE exposure, which also likely plays an important role in the effects observed. Investigation of these variables in future studies would provide a more comprehensive understanding of how our aCIE exposure, as well as a more protracted abstinence period following adolescent alcohol exposure, influences glutamate transmission in the male BLA.

Interestingly, our sEPSC findings in our female animals are somewhat consistent with these previous works, as we found increased glutamate release in aCIE females and a trend toward increased postsynaptic effects. A study by Morales et al. found that females exhibited pre- and postsynaptic changes to the glutamate system following a 10-day exposure and acute withdrawal (Morales et al., 2018). Interestingly, females were more resistant to adolescent alcohol exposure-induced alterations to the glutamate system, in that females required a full 10-day exposure before exhibiting the changes described above, whereas males required only a 7-day exposure to begin exhibiting alterations (Morales et al., 2018). In contrast to these findings in the BLA, including the current study, in the BNST of females sEPSC frequency was enhanced and NMDA receptor-mediated long term depression was attenuated during acute withdrawal, with this effect being absent 30 days later (Carzoli et al., 2019; Kasten et al., 2020). Given that few studies have examined physiological alterations following adolescent alcohol exposure in females, these studies along with our findings point toward important sex-dependent adaptations in neurotransmission in anxiety-related brain areas, particularly the BLA, which may underlie, in part, some of the well-described sex differences in behavior associated with adolescent alcohol exposure.

In contrast to the glutamate system, we did not find aCIE-induced alterations to GABA transmission in either males or females. Although long-term alterations to inhibitory transmission in the BLA following adolescent alcohol exposure have not been examined, acute withdrawal from chronic intermittent ethanol exposure in presumptive (based on weight) adolescent males showed that presynaptic GABA release was suppressed 24 hours into abstinence, and this was specific to feed-forward inhibitory synapses that mediate top-down regulation of BLA excitability (Diaz et al., 2011). In this same study, local feedback inhibitory synapses did not exhibit this decrease in GABA release, indicating that alterations to inhibitory transmission are synapse-dependent. Postsynaptic alterations to the GABA system at both synapse types were also found during acute withdrawal (24 hours into forced abstinence), although this appeared to differentially affect specific GABA_A_ receptor subunits at different synapses (Diaz et al., 2011). Importantly, these synapses are localized on different compartments on pyramidal cells within the BLA, with feedforward inhibitory synapses located along the external capsule and synapse onto distal dendrites, while local feedback inhibitory synapses are spread throughout the BLA and synapse onto proximal dendrites (Marowsky, Yanagawa, Obata, & Vogt, 2005; Silberman, Ariwodola, & Weiner, 2009; Woodruff & Sah, 2007). Because of the location of these different synapses onto pyramidal cells, sIPSCs are most likely derived from GABA release from local interneurons (Silberman et al., 2009). In the present study, our sIPSC recordings were therefore likely sampled from local feedback inhibitory synapses. Based on this, our finding that aCIE did not alter sIPSC frequency in males is potentially consistent with that from Diaz and colleagues. Additionally, since we did not observe changes in sIPSC amplitude, it appears that aCIE does not produce long-term postsynaptic changes to GABA transmission; however, we did not examine any receptor- or subunit-specific mechanisms so it is difficult to determine how our findings relate to previous studies. As with our glutamate results, future exploration of these and other potential mechanisms would provide a more thorough understanding of how our aCIE paradigm and protracted abstinence may influence this system. The present study is also the only study to examine adolescent alcohol exposure-induced alterations to GABA transmission in the BLA of females. While our GABA findings from males and females were similar, differences in BLA physiology have previously been found between males and females (Blume et al., 2017), so further exploration of this system in females is necessary.

We also assessed whether aCIE exposure would alter BLA KOR function. In control males, we found that KOR activation did not change GABA or glutamate transmission, a finding that is consistent with previous work from our lab (Przybysz et al., 2017). Interestingly, in aCIE males, KOR activation significantly reduced sIPSC frequency. The aCIE exposure also shifted the number of individual cells that responded to the agonist toward greater overall inhibition. aCIE exposure also changed the way KOR activation modulated glutamate transmission, with no effects in control males, but a significant reduction from baseline in sEPSC amplitude. These data indicate that aCIE exposure disrupts BLA KOR function in males by suppressing both the GABA and glutamate systems. Interestingly, although aCIE shifted the proportion of cells that responded to the KOR agonist in females, on average, this did not produce significant changes in GABA or glutamate transmission in either female group. While this is the first study to examine the impact of adolescent ethanol exposure on KOR modulation of synaptic function in the BLA, several studies have investigated the effects of chronic ethanol exposure on the KOR system in adulthood in various other brain regions including the extended amygdala. One study using repeated binge-level administration of ethanol found that KOR mRNA was increased relative to controls in tissue containing both BLA and central amygdala (D’Addario et al., 2013). Increased dynorphin-A immunoreactivity and dynorphin-A stimulated G-protein receptor coupling were also found in the central amygdala following induction of ethanol dependence and acute withdrawal (Kissler et al., 2014). Consistent with findings in the amygdala, KOR mRNA expression was also increased in the BNST during acute withdrawal from chronic ethanol exposure (Erikson et al., 2018). Finally, withdrawal from chronic ethanol exposure increases KOR regulation of dopamine transmission in the nucleus accumbens which may underlie reduced dopamine release in ethanol-dependent rats (Karkhanis et al., 2016; Rose et al., 2016). Together, these studies demonstrate that chronic ethanol exposure in adulthood generally causes an upregulation and/or hyperfunction of the KOR system in the extended amygdala. While we did not directly examine KOR expression in the present study, our results are consistent with increased expression and/or function of the KOR system as U69593 produced significant reductions in glutamate and GABA in aCIE males, an effect that was absent in control males in both the current and our previous study (Przybysz et al., 2017).

Notably, the present study differs from the work described above in that our electrophysiological testing occurred after a protracted abstinence period, rather than during acute withdrawal. Studies examining changes to *expression* of the KOR system have found that dynorphin-B peptide levels were upregulated 21 days following adolescent alcohol exposure (Granholm, Segerstrom, & Nylander, 2018; Lindholm, Ploj, Franck, & Nylander, 2000), but current studies examining long-term changes to KOR *function* following chronic alcohol exposure are limited. One study examining the role of the KOR system in the alcohol deprivation effect following 18 days of abstinence found that the selective KOR antagonist JDTic reduced the reinstatement of alcohol intake (Uhari-Väänänen et al., 2019), while another found that the KOR system was still actively involved in mediating abstinence-induced stress responsiveness six weeks after chronic alcohol exposure (Gillett, Harshberger, & Valdez, 2013). Another study specifically examining the role of central amygdala KORs in alcohol intake during acute and protracted withdrawal found that infusion of a KOR antagonist into the central amygdala reduced alcohol consumption during acute withdrawal, and that this effect lasted throughout protracted abstinence (Kissler & Walker, 2016). These studies together indicate that the KOR system is both altered by chronic alcohol exposure in adulthood and remains in this altered state for up to six weeks following termination of exposure. Our results compliment this work in that we show that at least 30 days following termination of aCIE exposure, male KOR function is disrupted. We also expand upon this work by revealing these effects in a brain region that has not previously been studied in this context. Importantly, all of the studies described above used an alcohol exposure paradigm that took place during adulthood, while the present work used an exposure during adolescence. The BLA is heavily involved in behaviors associated with the long-term effects of both adult and adolescent alcohol exposure including anxiety, stress responsiveness, and reward processing, so this work opens a broad avenue for future investigation into the behavioral implications of KOR dysfunction following alcohol exposure at multiple developmental ages.

Work investigating the role of the KOR system in females is much less abundant than work using males, but studies including both sexes have consistently demonstrated that females may be less sensitive to KOR manipulation [for review, see (Chartoff & Mavrikaki, 2015)]. Notably, this has been shown in behaviors related to affective states such as anxiety, social interaction, depression, and reward sensitivity (Przybysz, Varlinskaya, & Diaz, 2020; Russell et al., 2014; Varlinskaya, Spear, & Diaz, 2018), all of which engage the BLA in some capacity. It is therefore interesting that we did not observe differences in how KOR activation modulates BLA neurotransmission between control males and females. On the other hand, in aCIE-exposed animals, we found that KOR activation altered neurotransmission in males but had no effect in females. While, to our knowledge, there are no other studies examining long-term effects of alcohol exposure on KOR function in females, this finding is consistent with work showing that females are resistant to acute KOR-mediated responses to manipulations like stress. For example, we found that females with a history of restraint stress were less sensitive to the socially-inhibiting effects of KOR activation (Varlinskaya et al., 2018). We also recently showed that social behavior in either adolescent or adult females was not affected by forced swim stress, a manipulation that has been shown to engage the KOR system (Varlinskaya et al., 2020). While the present study provides a first look at how long-term abstinence from adolescent alcohol exposure may disrupt KOR function in females, it is clear that additional work is needed to fully understand these effects across the brain and their influence on sexspecific behavioral profiles.

Another important distinction between the present study and previous work examining the effects of adolescent alcohol exposure is that we used a moderate-level exposure paradigm. Adolescents who consume alcohol often consume it in a binge-like pattern (Spear, 2014), but according to the National Drug Use and Health report from 2018, a similar percentage of adolescents consume alcohol in amounts that do not reach binge levels (Substance Abuse and Mental Health Services Administration, 2018). Despite this, the majority of preclinical work dedicated to understanding the neural consequences of adolescent alcohol exposure has used binge-level exposure, with very few studies explicitly examining the effects of low-moderate exposure. One study that ran a battery of behavioral assays following 4-week voluntary intermittent alcohol consumption, an exposure that likely yielded more moderate-level BECs, found no differences in novel object recognition or anxiety-like behavior two weeks following the final exposure (Sanchez-Marin et al., 2020). Interestingly, however, this study did identify several alterations to endocannabinoid- and neuroinflammation-related mRNA levels in discrete brain regions, indicating that a lower-level adolescent exposure may still produce persistent neural alterations which may require an additional stimulus to illicit an observable behavioral alteration. This is supported in the present work, and highlights the importance of examining the long-term neural consequences following low-moderate adolescent alcohol exposure.

Overall, the current study found that moderate aCIE exposure followed by protracted abstinence enhanced BLA glutamate transmission and produced functional alterations to the BLA KOR system, and that these effects were sex dependent. This study is the first to demonstrate that adolescent alcohol exposure produces long-term changes to BLA physiology and provides further evidence of the vulnerability of the KOR system to insults during developmentally sensitive periods of the lifespan [for review, see (Diaz et al., 2017)]. There are numerous additional questions that are raised by the findings presented here which open several broad areas of further investigation. Specific questions that arise from this work include examination of the mechanisms underlying potential changes to inhibitory and excitatory neurotransmission within the BLA and the behavioral consequences of disrupted basal and KOR-mediated BLA physiology. Additionally, this work highlights a large gap in our overall understanding of how developmental alcohol exposure may disrupt KOR function throughout the brain, as well as the long-term physiological dysfunction within the BLA and throughout the brain following exposure to various levels of alcohol during adolescence. Finally, this study demonstrated that there are important sex differences in the neural consequences of moderate adolescent alcohol exposure, further emphasizing the need to include females in many of these explorations. Overall, advancing our understanding of how adolescent alcohol exposure produces alterations to the excitatory/inhibitory balance in the brain and the systems that modulate it, many of which may confer vulnerability to affective dysfunction and alcohol use disorder in adulthood, will be critical to developing better therapeutic tools to better treat these disorders across the population.

